# Highly Realistic Whole Transcriptome Synthesis through Generative Adversarial Networks

**DOI:** 10.1101/2022.11.10.515980

**Authors:** Suneng Fu

## Abstract

The transcriptome is the most extensive and standardized among all biological data, but its lack of inherent structure impedes the application of deep learning tools. This study resolves the neighborhood relationship of protein-coding genes through uniform manifold approximation and projection (UMAP) of high-quality gene expression data. The resultant transcriptome image is conducive to classification tasks and generative learning. Convolutional neural networks (CNNs) trained with full or partial transcriptome images differentiate normal versus lung squamous cell carcinoma (LUSC) and LUSC versus lung adenocarcinoma (LUAD) with over 96% accuracy, comparable to XGBoost. Meanwhile, the generative adversarial network (GAN) model trained with 93 TcgaTargetGtex transcriptome classes synthesizes highly realistic and diverse tissue/cancer-specific transcriptome images. Comparative analysis of GAN-synthesized LUSC and LUAD transcriptome images show selective retention and enhancement of epithelial identity gene expression in the LUSC transcriptome. Further analyses of synthetic LUSC transcriptomes identify a novel role for mitochondria electron transport complex I expression in LUSC stratification and prognosis. In summary, this study provides an intuitive transcriptome embedding compatible with generative deep learning and realistic transcriptome synthesis.

**Significance Statement:** Deep learning is most successful when the subject is structured. This study provides a novel way of converting unstructured gene expression lists to 2D-structured transcriptome portraits that are intuitive and compatible with a generative adversarial network (GAN)-based deep learning. The StyleGAN generator trained with transcriptome portrait libraries synthesizes tissue- and disease-specific transcriptomes with significant diversity. Detailed analyses of the synthetic transcriptomes reveal selective enhancement of clinically significant features not apparent in the original transcriptome. Therefore, transcriptome-image-based generative learning may become a significant source of de novo insight generation.

## INTRODUCTION

Transcriptional regulation enables the specification of cell fate, physiological response, and disease pathogenesis. Since the advent of microarray and RNA sequencing technologies, transcriptome profiles have become the most comprehensive among all omics data (Lashkari et al., 1997; Nagalakshmi et al., 2008; Wilhelm et al., 2008). However, machine learning tools with transcriptomic data remain limited and have yet to transform the field of biology (Eraslan et al., 2019).

Typically, machine learning is most successful when dealing with structured data, like images and languages. The recent success in protein structure modeling with AlphaFold and Rosetta are also examples of machine learning with structured, ordered data (Anishchenko et al., 2021; Jumper et al., 2021). Significant efforts have been made to capture the power of computer vision-oriented tools to transform transcriptome learning by converting gene expression lists to structured images. For example, Lyu and Haque first used the chromosome location of genes as coordinates to convert the one-dimensional transcriptome profile to two-dimensional images (Lyu and Haque, 2018), which was replicated in more recent efforts by Chen and colleagues (Chen et al., 2021). Sharma and colleagues pioneered converting various kinds of non-image/tabular data to images with DeepInsight, and they were particularly successful (99% accuracy) in embedding and classifying RNAseq data with t-distributed stochastic neighbor embedding (t-SNE) and kernel principal component analysis (kPCA) techniques (Sharma et al., 2019). Bazgir and colleagues used the Bayesian multidimensional scaling approach in REFIND (Bazgir et al., 2020). Although all of them were moderately successful in sample classification, the images were not intuitive, nor did they demonstrate new learning capabilities.

Here, I embedded the mouse and human transcriptome as a phoenix-like object, thus named phoenix transformation, and the resulting transcriptome image demonstrated generative capacities for realistic transcriptome synthesis.

## RESULTS

### Transcriptome structurization through manifold embedding

Consider all genes in a genome as set X, and they interact with each other on an n-dimensional Riemannian space; we have X × X → ℝ^n^ (Fig. 1a). Through dimension reduction techniques, all genes in a genome may be mapped onto a two-dimensional Cartesian space. This study chose the uniform manifold approximation and projection (UMAP) (McInnes et al.) algorithm for dimension reduction. Three expression data sets covering the whole life span of mice were analyzed by UMAP: DRA000484 for zygotes and fertilization from the DBTMEE project (Park et al., 2015), ENCSR574CRQ for mouse early embryogenesis from the ENCODE3 project (He et al., 2020), and GSE132040 for mouse aging from the Tabula Muris Senis consortium (Almanzar et al., 2020) (Fig. 1b; Supplementary Table 1). The resultant Cartesian coordinates (Supplementary Table 2) were rotated and stretched through sequential multiplications of 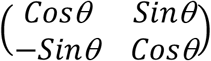 and 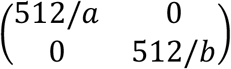 to produce pixel map coordinates for 512×512 transcriptome images (Fig. 1b).

**Figure 1.**
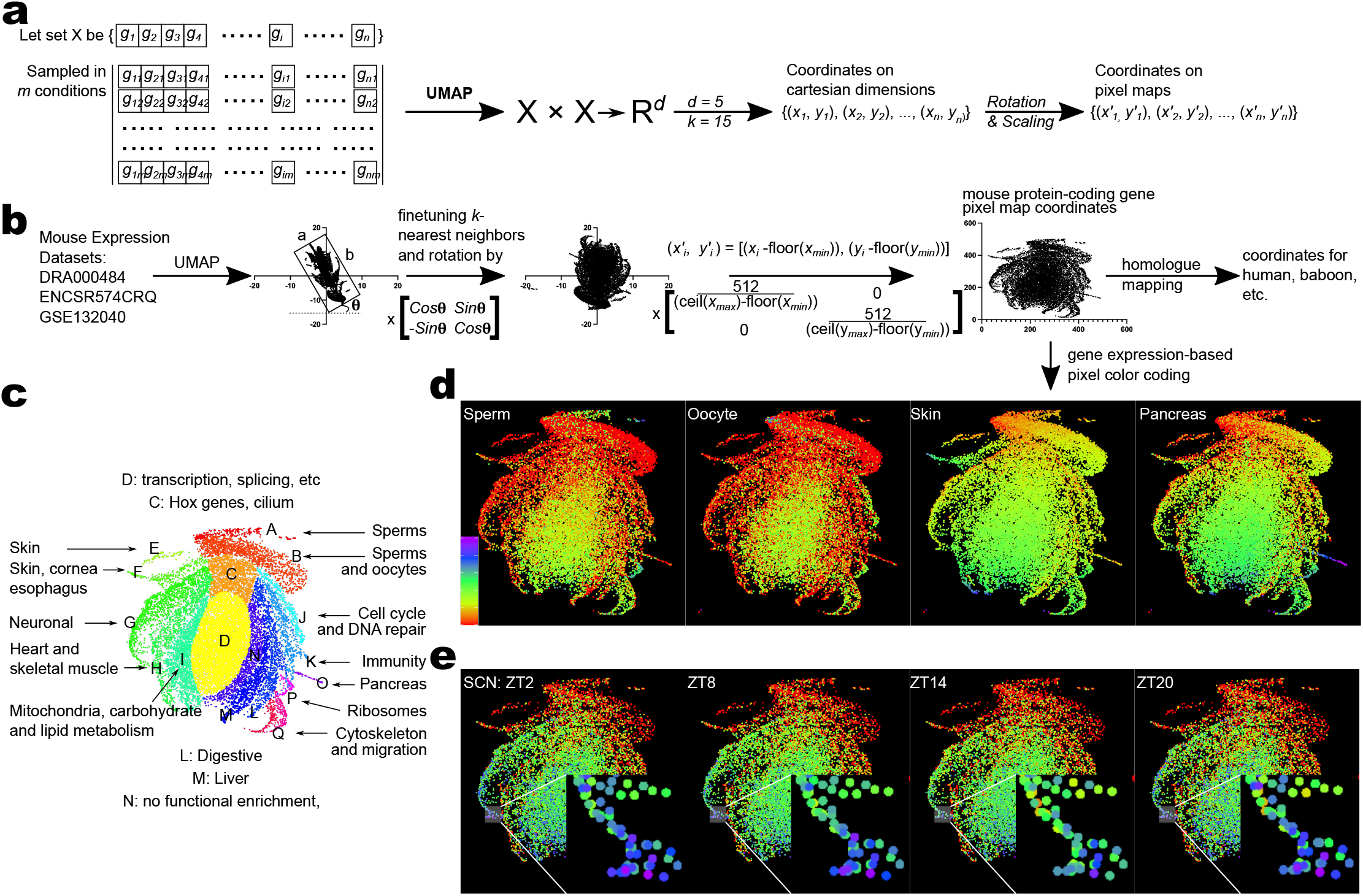
Uniform manifold approximation and projection (UMAP) of the transcriptome. **(a-b)**, Theoretical (**a**) and practical (**b**) pipelines of phoenix transformation for 2-D transcriptome visualization. Protein-coding genes in the mouse genome (set X) is mapped onto a Riemannian space of 5 dimensions with expression dataset (DRA000484, ENCSR574CRQ, GSE132040, combined in Supplementary Table S1). The cartesian coordinates of the first two dimensions are converted to pixel map coordinates (Supplementary Table S2). Pixel map coordinates for protein-coding genes in human and baboon genome are mapped through their mouse homologues (Supplementary Table S3, 4). **c**, Compartmentalization of the transcriptome manifold. Each compartment is color-coded based on their enrichment for tissue-specific genes or gene ontologies (GO). Gene compartment assignment is provided in Supplementary Table S5. Gene set enrichment for each compartment is provided in Supplementary Table S6. **d**, Representative mouse tissue transcriptome images. Individual genes are represented as dots rather than singular pixels and in ascending orders of expression. The fragments per kilobase of exon per million reads (FPKM) expression of each gene was log2 transformed then multiplied by 14, clipped between {1, 255}. Color scales from 0 (red) to 255 (purple). Corresponding gene expression data are provided in Supplementary Table S7. **e**, Representative baboon circadian transcriptome from the suprachiasmatic nucleus (SCN). Inserts show zoom-in views of a section of neuron-specific gene expression oscillation across circadian time at 3072×3072 pixel resolution. ZT, zeitgeber time; SCN, suprachiasmic nucleus. Gene expression data are provided in Supplementary Table S8.

The gradient 6, width a, and length b describe the minimal rectangle that encircles all coordinates (Fig. 1b), all of which were determined similarly as in DeepInsight (Sharma *et al*., 2019). The pixel coordinates were then used for color-coding of gene expression information and the production of transcriptome images. Please refer to Fig. 1 and Methods for detailed information about computing and rendering the transcriptome image.

Together, 20,424 protein-coding genes in the mouse genome were mapped onto 18,545 unique coordinate pairs (Fig. 1b). The coordinates for protein-coding genes in human and baboon genomes were determined through homolog mapping (Fig. 1b). Fig. 1c shows the mouse transcriptome resolved as a phoenix-shaped manifold; tissue-specific and function-related genes congregate to form body parts of the phoenix. Genes located in the “head” (zone A) are sperm-specific, and genes located in the “neck” (zone B) are selectively expressed in reproductive cells/tissues, including sperms, oocytes, and the testis (Fig. 1d; Supplementary Fig. 1a). Underneath the neck is zone C that enriches for body plan genes expressed during early embryogenesis (Fig. 1c). Compartments connected to the body plan zone from the left are epithelial (E, F), neuronal (G), muscular (H), and respiratory/metabolic (I) genes. On the right are proliferation (J), immunity (K), and digestion-related (L, O) genes (Fig. 1c; Supplementary Fig. S1a). Underneath zone C, there resides a large group of poorly-resolved housekeeping genes (D, M, N, P, Q; Fig. 1c).

Despite compartment-based clustering of tissue-specific genes, their expression level varies across circadian time. For example, the neuronal zone gene expression in the baboon SCN transcriptome (Mure et al., 2018) image reaches a nadir at ZT14 (Fig. 1e), coinciding with SCN inactivity starting around 10 pm(Deboer et al., 2003). In contrast, non-neuronal tissues may elevate neuronal gene expression because of heightened neuronal activation of the target tissue, in this case, the lung at ZT0 (Supplementary Fig. S1b).

### Transcriptome image classification through convolutional neural networks

Cancer and normal tissue transcriptome images are visually distinct in the proliferation zone, with the cancer transcriptome exhibiting a blue shift in pixel coloration, i.e., higher levels of gene expression than normal (Fig. 2a). Additionally, the cancer transcriptome image often presents a variable degree of red-shift in tissue-specific zone coloration, i.e., reduced expression of normal tissue functions (Fig. 2a). For comprehensive transcriptome image classification, convolutional neural networks (CNNs) were trained and evaluated.

**Fig 2.**
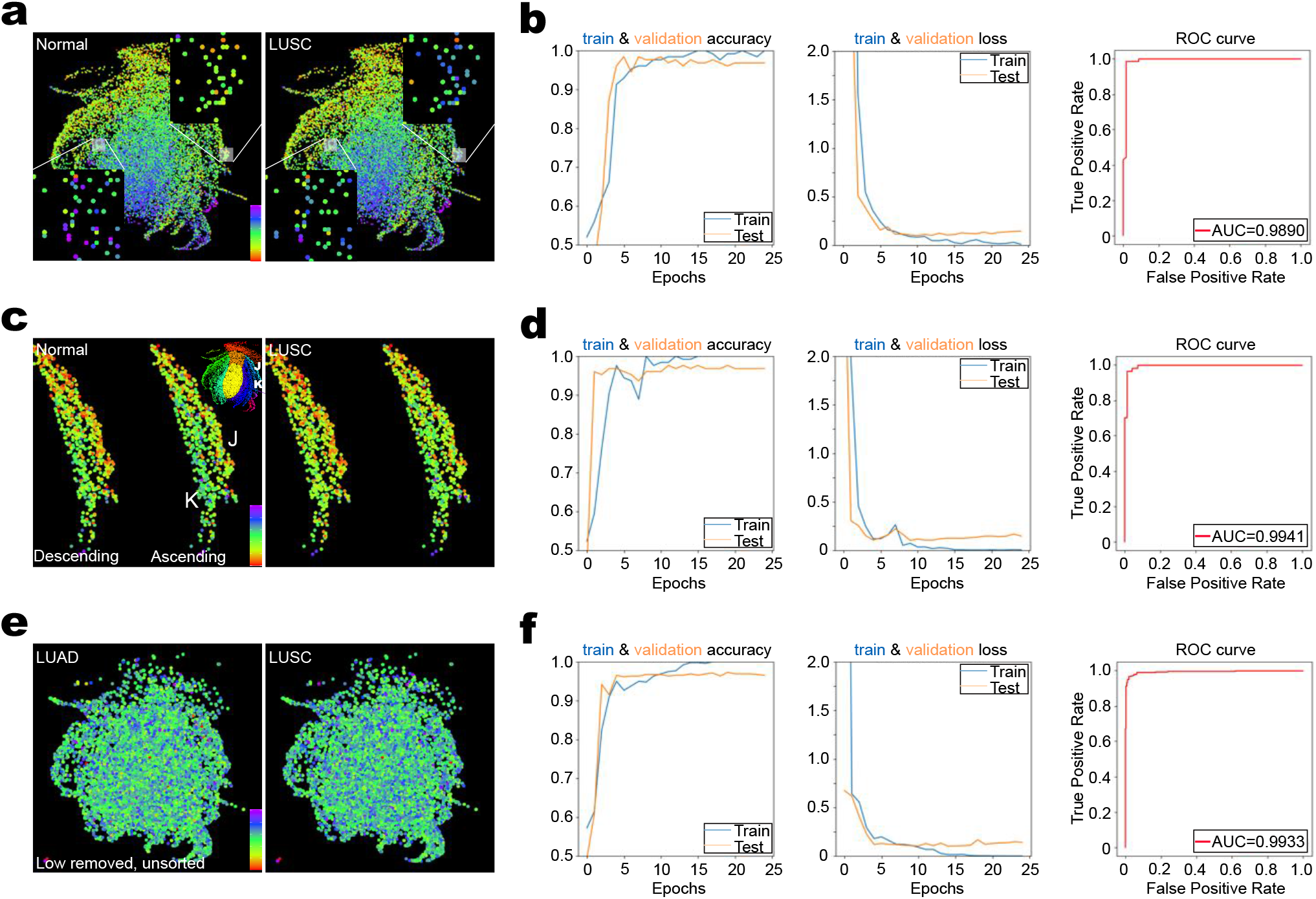
Lung cancer diagnosis and classification with transcriptome images. **a-b**, Images (**a**) and convolutional neural network (CNN)-based detection (**b**) of lung squamous cell carcinoma (LUSC) from cancer and neighboring normal tissue transcriptomes. Transcriptome image is rendered in ascending orders of gene expression and resized from 1024×1024 to 5122×512 pixels. Zoom-in views in (**a**) show upregulation of genes in the proliferation zone (upper right) and downregulation of lung-specific genes (lower left). Gene expression data for transcriptome classification is from GSE18842, 19804, and 27262 (combined in Supplementary Table S9). **c-d**, Images (**c**) and detection (**d**) of LUSC with images containing only genes in the proliferation zone (J) and immunity zone (K) from the same dataset as **a** and **b**. The left side of the image is rendered in descending orders of gene expression, and the right side is in ascending order. Image is rendered in 512×512 pixels. **e-f**, Images (**e**) and classification (**f**) of LUSC and LUAD transcriptomes with lowest quarter gene expression removed. The image is resized from 1024×1024 to 512×512 and not rendered according to the orders of gene expression. The data is from TCGA and normalized by edgeR (Supplementary Table S11).

First, the number of convolutional neural network layers needed to achieve effective transcriptome image classification was tested (Supplementary Fig. S2a). Supplementary Fig. S2b shows that two-layer CNNs quickly reached >90% accuracy for normal versus LUSC classification during the training process, followed by one-layer CNN. CNNs of 3∼5 layers failed to achieve effective classification in 20 epochs. The performance of Adam and stochastic gradient descent (Sgd) optimizers were also compared. The results showed the Adam optimizer to be more stable than Sgd during the training (Supplementary Fig, S2c, d). Therefore, two-layer CNNs with the Adam optimizer were chosen for transcriptome image classification.

Second, the type of transcriptome images suitable for classification was explored. The transcriptome image might be rendered and classified in a multitude of ways based on their level of gene expression, the inclusion of partial or whole transcriptomes, and image sizes. For cancer detection, CNNs trained with either complete transcriptome images rendered in the ascending orders of gene expression or partial transcriptome images comprised of only the proliferation and immunity genes achieved ∼ 98% of normal versus LUSC classification (Fig. 2a-d). The precision, sensitivity, and specificity level were around 96-98%, comparable with the state-of-the-art gene expression list-based XGBoost classifier (Table 1; Supplementary Fig. S2e) (Chen and Guestrin, 2016). However, cancer classification between LUSC and LUAD (lung adenocarcinoma) was less successful with full transcriptome images, but removing the bottom 25% of genes from the transcriptome images improved the model accuracy to ∼96% (Table 1; Fig. 2e, f). The sensitivity, specificity, and F1_scores were all around 94∼96% (Table 1). The XGBoost classifier achieved 100% accuracy with the same dataset (Table 1; Supplementary Fig. S2f). Therefore, the performance and the choice of transcriptome image best suited for classification are task-dependent.

**Table 1.**
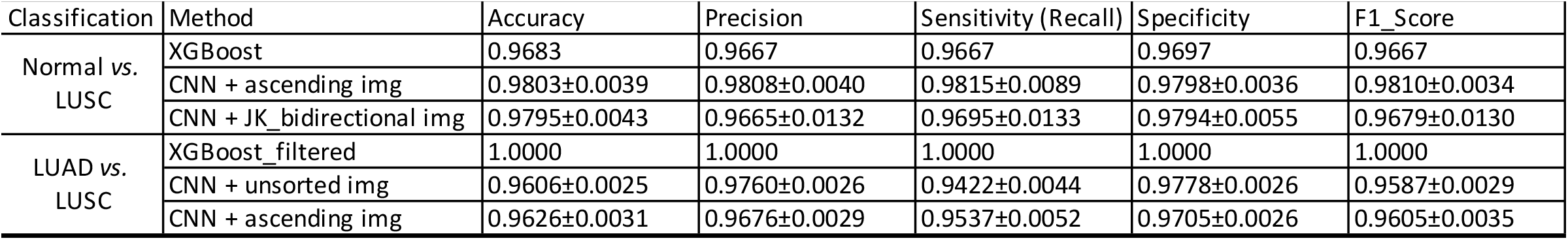

| Classification | Method | Accuracy | Precision | Sensitivity (Recall) | Specificity | F1_Score |
| --- | --- | --- | --- | --- | --- | --- |
| Normal vs. LUSC | XGBoost | 0.9683 | 0.9667 | 0.9667 | 0.9697 | 0.9667 |
|  | CNN + ascending img | 0.9803±0.0039 | 0.9808±0.0040 | 0.9815±0.0089 | 0.9798±0.0036 | 0.9810±0.0034 |
|  | CNN + JK_bidirectional img | 0.9795±0.0043 | 0.9665±0.0132 | 0.9695±0.0133 | 0.9794±0.0055 | 0.9679±0.0130 |
| LUAD vs. LUSC | XGBoost_filtered | 1.0000 | 1.0000 | 1.0000 | 1.0000 | 1.0000 |
|  | CNN + unsorted img | 0.9606±0.0025 | 0.9760±0.0026 | 0.9422±0.0044 | 0.9778±0.0026 | 0.9587±0.0029 |
|  | CNN + ascending img | 0.9626±0.0031 | 0.9676±0.0029 | 0.9537±0.0052 | 0.9705±0.0026 | 0.9605±0.0035 |

Lastly, I examined how gene positioning in the transcriptome image may impact image classification by scrambling their coordinates. Intriguingly, CNN models trained with transcriptome images with scrambled gene coordinates achieved >95% accuracy in LUSC versus LUAD classification (Supplementary Fig. S2g), comparable with models trained with unscrambled coordinates. These results suggest that transcriptome image classification depends on pixel-level information regardless of spatial positioning. Such an interpretation is consistent with the observation that CNN models of 1-2 layers outperformed those with three or more layers (Supplementary Fig. S2b).

### Generative adversarial network training and realistic transcriptome synthesis

Unlike gene expression lists, images are suitable for training with generative adversarial networks (GAN) that can learn rules of play from the ground up (Goodfellow et al., 2014). However, the transcriptome portrait is a dot plot unlike conventional images used for GAN training. Therefore, I asked whether classical generative models may be trained to synthesize transcriptome images and used for further analysis (Fig. 3a). The StyleGAN2-ADA framework (Karras et al., 2020) is the best performing model in the field in synthesizing highly realistic pictures, thus is chosen for this study (Fig. 3b).

**Figure 3.**
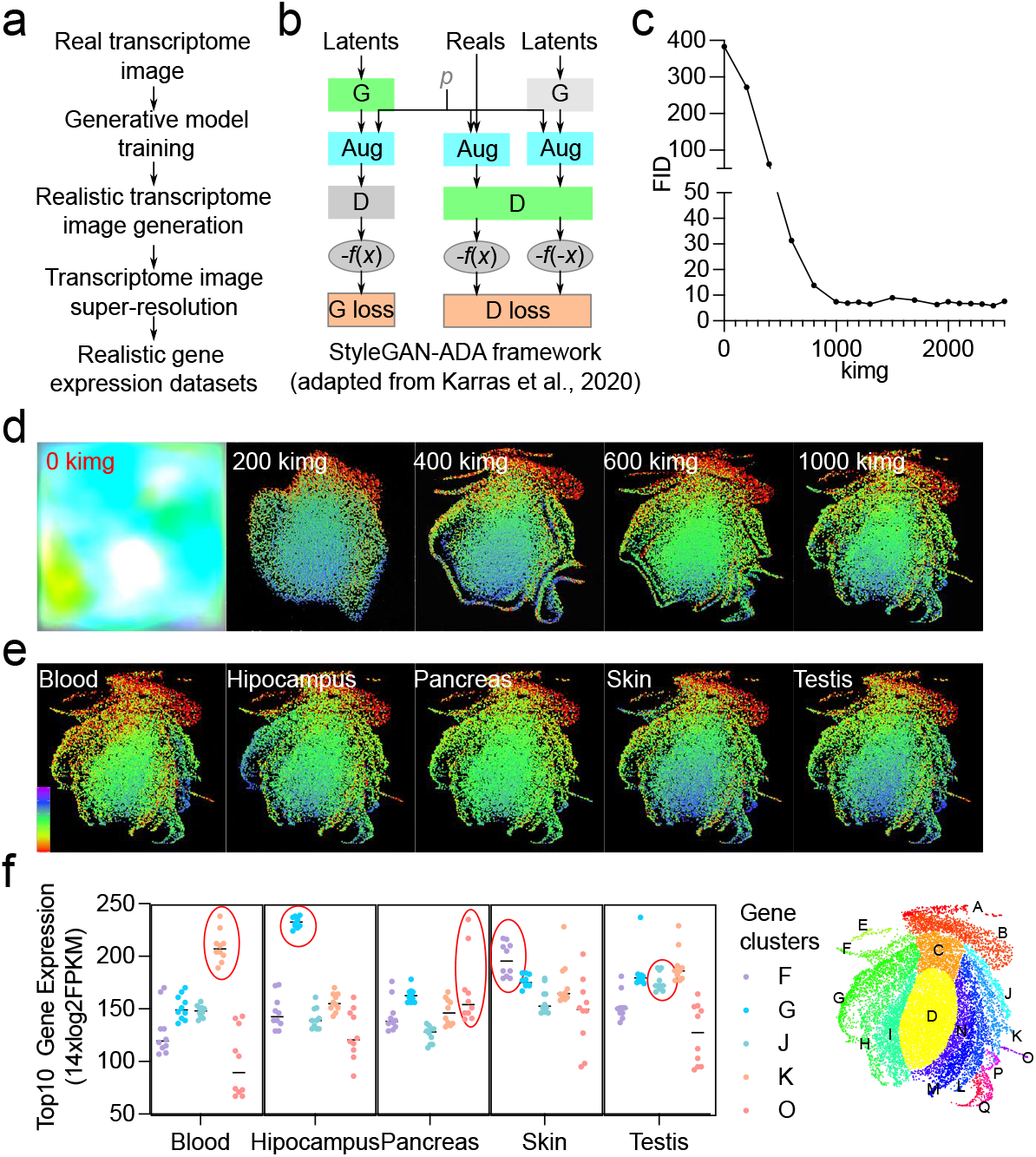
Generation of tissue-specific transcriptomes with generative adversarial networks (GAN). **a**, Illustration of the pipeline for transcriptome image generation and gene expression list retrieval. **b**, Illustration of the StyleGAN2-ADA network. The generative (G) and the discrimating (D) network competes in fooling and revealing each other evaluated by the G- and D-loss scores. **c**, The progression of StyleGAN2-ADA conditional network training with 93 classes of normal and cancer transcriptome images. The Fréchet inception distance (FID) measures the overall similarity between the real (training image) and generated image space. The TcgaTargetGtex dataset is used for training (Supplementary Table S13) **d**, Examples of transcriptome image produced by the generative network during the training process. **e**, Generatiion of tissue-specific transcriptome image by the final generative network. **f**, The level of top10-expressed genes from F, G, J, K, O clusters characteristic of skin, hipocampus, testis, blood, and pancreas tissues, respectively, in the GAN-synthesized transcriptomes. Reconstructed gene expression dataset is provided in Supplementary Table S15.

Four different transcriptome image renditions were tested for their suitability for StyleGAN training: 1) 3072×3072 transcriptome images, 2) 1024×1024 transcriptome images rendered in ascending orders of gene expression, 3) 1024×1024 transcriptome images rendered according to the alphabetic orders of gene symbols, and 4) the same transcriptome image as #2 but with scrambled gene coordinates. All images were resized to 512×512 and referred to as 3072×3072, ascending, and unsorted, respectively. Each training library contained 9,300 transcriptome images evenly sampled from 93 tissue/cancer classes from the TcgaTargetGtex data set. The training progress was monitored by the Fréchet Inception Distance (FID) between actual and GAN-generated images(Martin Heusel et al., 2018). Because of difficulties associated with deconvoluting low-resolution images into gene expression lists, original transcriptome images rendered at 512×512 resolutions were not tested for StyleGAN training.

The training results show that StyleGAN is more stringent in image specifications than CNN. Among all four image configurations, the conditional StyleGAN2-Ada network trained with ascending transcriptome images achieved the best FID score (17.88) without parameter finetuning (Supplementary Fig. S3a). Scrambling gene coordinates impaired StyleGAN training, and the best FID score achieved during training was 106.37 (Supplementary Fig. S3a). Training with the unsorted transcriptome image library caused an inversion of the background from black to white (Supplementary Fig. S3b). Meanwhile, training with 3072×3072 images led to a loss of the head region (Supplementary Fig. S3c). The impact of batch size on training was also assessed, and it was found that a batch size of 16 overperformed batch sizes of 8 and 32 at default settings. Further finetuning of the batch 16 model reduced the FID score to 5.82 (Fig. 3c).

Besides the FID score, GAN-generated images were also visually evaluated during training. Fig. 3d shows that the StyleGAN generator learned the overall shape of the transcriptome embedding within 200 thousand images (kimg) iterations, and it continued to refine the segmentation upward to 1000 kimg (Fig. 3d). The final model produced transcriptome images with class-specific characteristics (Fig. 3e). For example, class 3 latent vectors synthesized transcriptome image with highest pixel coloration (blue to purple) in the immunity zone (K), characteristic of whole-blood transcriptomes. Class 18 latent vector produced images with the highest coloration in the neuronal zone (G), emulating the brain transcriptome. Pixel coloration in zone A and B of class 82 synthetic images and the zone O of class 66 synthetic images were not as high as their corresponding real-world testis and pancreas transcriptome images but higher than those in other tissues (Fig. 3e), indicating a partial retainment of tissue characteristics.

Super-resolution GANs (SRGANs) (Ledig et al., 2017) were trained to reduce pixel overlap between gene dots in the 512×512 images before converting the image to gene expression lists (Supplementary Fig. S3d). As the training progressed, the generator loss (G_loss, Supplementary Fig. S3e), a metric for assessing how well the generator fooled the discriminator, rapidly decreased. Meanwhile, the peak noise-to-signal ratio (PNSR) of the produced image steadily increased to a plateau (Supplementary Fig. S3f). Visually, the number of empty circles in the transcriptome image progressively decreased (Supplementary Fig. S3g). Notably, the image file size also decreased during training (Supplementary Fig. S3g), consistent with the reduction of superfluous ring-like structures in the transcriptome image. After converting back to gene expression lists, expression values from the super-resolution transcriptome image correlated with their original expression values with an R^2^ value of 0.9541 and a slope of 0.9943 (Supplementary Fig. S3h). The median absolute error was 0.43 log_2_CPM(Supplementary Fig. S3i).

The performance of SRGANs in the super-resolution of StyleGAN-generated transcriptome images lagged considerably compared to real low-resolution pictures (see the right-most panel in Supplementary Fig. S3g). Still, the gene expression lists from the StyleGAN-synthesized tissue-specific transcriptome images retained their respective, actual tissue transcriptome signatures. For example, the median expression level of the top10 genes deduced from the immunity (K) zone of the StyleGAN-synthesized whole-blood transcriptome image was ∼16X higher than in other tissues (Fig. 3f). Similarly, the expression values of top10 neuronal genes (cluster G) from the synthetic hippocampus transcriptome approached 250 (14xlog_2_CPM), far exceeding neuronal gene expression in other tissues’ synthetic transcriptomes (Fig. 3f).

### Feature retention and analysis of StyleGAN-generated lung cancer transcriptomes

The generator synthesized one hundred transcriptome images each for LUAD (class 57) and LUSC (class 58) with (*φ* = 0.7) or without (*φ* = 1.0) latent space truncation to systematically evaluate the fidelity, diversity, and utility of GAN-generated transcriptomes. Truncation of the latent space (*φ* < 1) was reported to improve image quality(Andrew Brock et al., 2019).

Fig. 4a shows notable variations in the epithelial (zone F) and immunity (zone K) compartment pixel coloration from GAN-synthesized LUAD and LUSC transcriptome image samples. After conversion to numeric gene expression lists, the number of highly expressed genes in the GAN-synthesized transcriptome was notably lower than the real transcriptome (Fig. 4b), consistent with signal loss observed during the super-resolution process, especially for the GAN-generated images (Supplementary Fig. S3g). Still, the expression levels of top20 genes from the epithelial zone changed from seed to seed in both LUAD and LUSC synthetic transcriptomes (Fig. 4c), confirming diversity among the synthetic transcriptomes. Truncation of the latent space with a *φ* = 0.7 lowered epithelial gene expression levels but retained between-sample variations.

**Figure 4.**
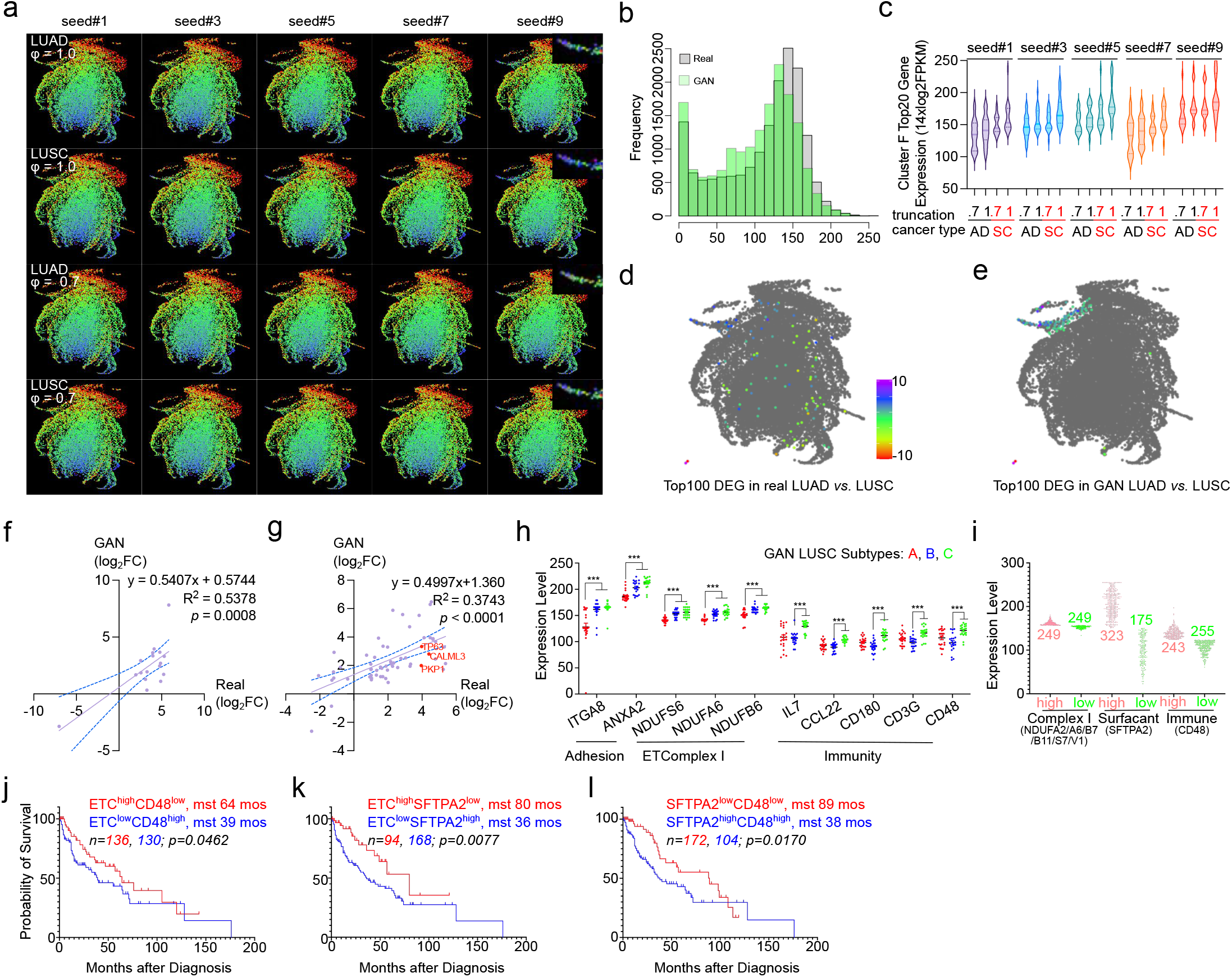
Synthesis and analysis of GAN-synthesized lung cancer transcriptome images. **a**, Realistic transscriptome images synthesized by trained StyleGAN2-ADA generative networks with seed 11, truncation 0.7, class=57 (LUAD) and 58(LUSC). b, Gene expression frequency distribution of real- and GAN-synthesized lung cancer transcriptomes. The real and GAN-reconstructed gene expression data are provided in Supplementary Table S16 and 17. c, The expression chracteristics of top20-expressed genes from cluster F in synthetic lung cancer transcriptomes. Corresponding reconstructed gene expression data are provided in Supplementary Table S18. **d-e**, The geometric distribution of top100 differentially-expressed genes (DEG) in the real (**d**) and GAN-synthesized (**e**) LUAD and LUSC transcriptomes. Color scale is log_2_-fold of change (FC). DEG expression data are provided in Supplementary Table S20, 22. **f**, Correlation of the 17 top100 DEGs from the real LUAD vs. LUSC transcriptome that are also differentially regualted in the GAN-synthesized LUAD and LUSC transcriptome. The *p-*value was calculated from linear regression test of whether the slope is significantly different from zero. **g**, Correlation of the 58 top100 DEGs from the GAN-synthesized LUAD vs. LUSC transcriptomes that are also regulated in the real LUAD vs. LUSC transcriptomes. The p-value was calculated as in (**f**). **h**, Differential expression of cell adhesion, mitochondria respiration complex I, and immunity genes in subtypes of GAN-synthesized LUSC transcriptome. The expression level is 14xlog_2_FPKM and provided in Supplementary Table S25. The *p*-value was calculated from two-tailed student *t*-test. **i**, The distribution of cell adhesion, complex I, and immunity gene expression in LUSC patient transcriptomes. **j-l**, Kaplan-Meier plots of patient survival after diagnosis with designated gene expression signatures. The *p*-value was calculated from log-rank test for between-group survival differences.

Regardless of truncation, epithelial gene expression in the synthetic LUSC transcriptome consistently exceeded those in the LUAD transcriptome generated by the same seed. Importantly, the hierarchical squamous epithelial structure is a unique pathological feature of LUSC tumor nests, and elevated expression of epithelial identity genes was implicated as a hallmark in differentiating LUSC from LUAD. Notably, all four upregulated genes from an eight-gene set used for differentiating LUSC from LUAD (Hamaneh and Yu, 2022) were epithelial identity genes located in zone F of the transcriptome image. StyleGAN training enhanced the epithelial identity of the LUSC transcriptome further. Fig. 4f shows that almost all DEGs between StyleGAN-synthesized LUAD and LUSC are concentrated in the epithelial zones E and F. For comparison, the top100 DEGs between real LUAD and LUSC transcriptomes are dispersed throughout the transcriptome pixel map. Further analyses revealed that only 17 of the top100 DEGs in the real transcriptome remained differentially regulated in the GAN-synthesized transcriptomes (Fig. 4g), while 62 of the top100 DEGs in the GAN transcriptome were DEGs in the real transcriptome (Fig. 4h).

Besides the comparative analysis of DEGs, I benchmarked the similarity between the actual and GAN-generated transcriptomes by examining how well models trained for the classification of real LUAD *versus* LUSC transcriptomes perform with GAN-synthesized ones. Supplementary Fig. 4a-d show that CNN models trained with full transcriptome images of real LUAD versus LUSC transcriptome images effectively differentiated GAN-synthesized ones. Specifically, the full-transcriptome image-based CNN achieved 91% accuracy in determining real LUSC and LUAD and 77% in GAN-synthesized image sets (Table 2). XGBoost classifier was trained after converting the real and GAN-synthesized transcriptome images back to numeric gene expression lists, and the XGBoost model trained with the real image-derived gene expression data achieved 91% accuracy in real LUAD versus LUSC classifications and 74% GAN-synthesized transcriptome classifications (Table 2; Supplementary Fig. S4e-h). Therefore, both CNN and XGBClassifier trained with real data was capable of classifying their corresponding synthetic transcriptome. The reduced efficacy might be attributed to the selective retention of LUSC epithelial identity and the loss of non-epithelial differences in the GAN-synthesized data.

**Table 2.**

| Method | Training + Testing | Accuracy | Precision | Sensitivity (Recall) | Specificity | F1_Score |
| --- | --- | --- | --- | --- | --- | --- |
| CNN* | Real for Real | 0.9090±0.0075 | 0.9238±0.0093 | 0.8972±0.0076 | 0.9217±0.0104 | 0.9101±0.0072 |
|  | Real for Synthetic | 0.7692±0.0074 | 0.8726±0.0157 | 0.6406±0.0342 | 0.8979±0.0200 | 0.7308±0.0160 |
| XGB** | Real for Real | 0.9100 | 0.8846 | 0.9388 | 0.8824 | 0.9109 |
|  | Real for Synthetic | 0.7400 | 0.6716 | 0.9184 | 0.5686 | 0.7759 |

\*CNN model was trained with full transcriptome images of LUAD and LUSC rendered in ascending orders of gene expression.
\*\*XGBClassifier was trained with top40 hallmark genes calculated by edgeR and validated by random forest in ranger from tabular expression data reconstructed from LUAD and LUSC transcriptome images.

The phenomenon of selective feature enhancement/elimination during StyleGAN training extended into the subtype structures of LUSC. Hierarchical cluster analysis of the synthetic LUSC transcriptomes showed a two-step bifurcation into three major subsets (Fig. 4h; Supplementary Fig. S4i). Gene set enrichment analysis (GSEA) for DEGs between clusters I and II showed an over-representation of cell adherence, mitochondria electron transport complex I, and EGFR signaling functions. Meanwhile, DEGs between clusters IIa and IIb were enriched for membrane proteins and immune response functions. For comparison, DEGs at both tier-I and tier-II branch points of real LUSC transcriptome subtypes were enriched for immunity genes but not mitochondria electron transport complexes (see Supplementary Tables associated with Supplementary Fig. S4i).

Significantly, the novel mitochondria signature enhanced during StyleGAN training provided critical prognosis values for LUSC patients. For example, although the expression levels of ETC I and immune response gene CD48 alone did not predict survival (Supplementary Fig. S4j, k), patients with a combined high ETC I and low CD48 expression (ETC^high^CD48^low^) had a median survival time (MST) of 64 months versus 39 months for patients of ETC^low^CD48^high^ (Fig. 4j). Combining ETC^high^ or CD48^low^ with another prognosis marker SFTPA2^low^ extended MST further to 80 (Fig. 4k) and 89 months (Fig. 4l), respectively. SFTPA2^low^ alone extended MST to 70 months in LUSC patients but reduced LUAD patients’ survival to 45 months after diagnosis (Supplementary Fig. S4l, m).

## DISCUSSION

In summary, this study demonstrates the utility of a phoenix-like 2D transformation of transcriptome data in sample classification and generative learning. Although gene coordinates seem insignificant for transcriptome image classification, it is crucial for the successful training of the StyleGAN network, in which the loss function is not driven by pixel intensity, *per se*, but by the distribution patterns of pixel intensities. Moreover, the coordinate system provides an intuitive way of visualizing tissue identity and gene set functional enrichment.

Despite this study’s proof-of-concept nature, CNN models trained with transcriptome images achieved comparable performance in sample classification with the state-of-the-art XGBoost algorithm. A combination of CNN models trained with different renditions of transcriptome pictures may even exceed the performance of XGBoost under challenging circumstances. Additionally, the preferential feature preservation/enhancement during StyleGAN training provides novel insights into the transcriptome not afforded by expression list-based machine learning models. An in-depth analysis of the StyleGAN latent space shall reveal the signatures of each of the 93 classes of normal and cancer tissues and valuable cancer-specific drug targets. Through transfer learning, StyleGAN and other generative adversarial networks may learn disease mechanisms with limited samples, e.g., rare diseases, patient-specific pathologies, highly infectious agents not amenable to large-scale experimentations, *etc*. Because the transcriptome image is essentially a dot plot distinct from real-life photos, extensive experimentation is needed to develop best practices for StyleGAN training. With additional technical refinement, realistic transcriptomes and their latent space shall be broadly valuable for biomedical research in the near future.

## Methods

### Tools

The following tools were used for phoenix transformation and subsequent analyses: R Studio, PyCharm, Prism 9, SRGAN, StyleGAN2-ADA, DAVID web server, and the STRING web server.

### Data

Mouse gene coordinates were calculated from GSE132040, ENCSR574CRQ, and DRA000484. The downloaded RNAseq datasets were combined in Supplementary Table 1. The baboon dataset used for visualization of transcriptome change in circadian was GSE98965 and provided in Supplementary Table 6. The LUSC dataset (GSE18842, 19804, 27262) used for visualization and training was downloaded from CuMiDa and provided in Supplementary Table 9. The TcgaTargetGtex dataset used for generative adversarial network training was downloaded from https://toil-xena-hub.s3.us-east1.amazonaws.com/download/TcgaTargetGtex_RSEM_Hugo_norm_count.gz and provided in Supplementary Table 12.

### Projection of mouse protein-coding genes onto a two-dimensional space

The Cartesian coordinates of protein-coding genes in the genome were calculated by plot_umap (bio_plotr package) from three expression data sets: GSE132040, ENCSR574CRQ, and DRA000484 (Supplementary Table 1). The target Riemannian space dimension was set a 5, the number of the neighborhood was set at 15, and the distance matrix used Euclidian. The resulting coordinates (*x*_*i*_, *y*_*i*_) were rotated 30° by multiplication of 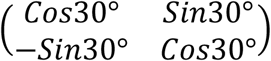, shifted the origin to (−17.5, -17.5), then stretched the x-y plane 16 fold by multiplying the new coordinates with 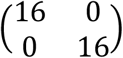. The final coordinates were rounded to integers for the eventual pixel map coordinates. Pixel map coordinates for protein-coding genes in other genomes were obtained from their mouse homologs. The final coordinates for mouse, human, and baboon genomes are provided in Supplementary Tables 2, 3, and 4.

### Transcriptome data transformation

For mouse and human RNAseq transcriptome datasets, the log2-transformed count per million (CPM) values were multiplied by 14, plus 1, rounded and clipped between [1, 255]. For the patient-paired normal versus LUSC microarray datasets, the transformation was performed through (n-3) * 1.7 * 14 + 1, rounded and clipped between [1, 255], where n was the probe signal from the microarray. For the baboon dataset, the transformation was round((log(n, 2)+3)*14), clipped between [1, 255], where n was the FPKM of individual genes. The transformed data is provided in Supplementary Tables 5-8.

### Tabular transcriptome data to image conversion

The transcriptome table was either sorted in ascending or descending orders of gene expression or arranged according to the alphabetic order of gene symbols. For genes with shared coordinates, only the data point rendered later in the gene expression table was preserved in the image. The transcriptome image was rendered in ggplot2 with each gene represented by a filled dot, positioned according to its pixel map coordinates calculated above, and color-coded by its expression levels. The x- and y-axis ranges were set at (0, 512), and the color scheme was a 5-color gradient. Pseudogenes with expression values of 0 and 255, pixel map coordinates of (20, 20) and (15, 15) were included in the transcriptome image for color uniformity. Each ggplot2 image were saved in three different sizes: 512×512 pixels (1.79 inches), 1024×1024 pixels (3.58 inches), and 3072×3072 pixels (10.74 inches).

For the construction of StyleGAN training libraries, the TcgaTargetGtex transcriptome dataset was assigned 93 classes based on tissue/cancer types. Each transcriptome class was randomly sampled either 500 times or 100 times to build libraries of 46,500 images and 9300 images. The transcriptome images were initially rendered at 1024×1024 pixels in ggplot2 and then resized to 512×512 pixels in ImageMagick.

For SRGAN training, each of the 93 TcgaTargetGtex classes was sampled 20 times to build three libraries of different image sizes: 512×512, 1024×1024, and 3072×3072. The 1024×1024 images were resized to 768×768 pixels for a fourth library. Transcriptome images from the LUSC class were deleted from the training library and reserved for quality evaluation.

### Model training

Initial model parameters for image classification were tuned by Keras Hyperparameter Tuner that compared 1-5 layers of multilayer perceptron (MLP) and convolutional neural networks (CNNs) with or without dropouts (https://keras.io/guides/keras_tuner/). The final CNN model consisted of two Conv2D layers of 32 and 64 filters, a kernel size of 3×3, and relu activation. Each was followed by a MaxPooling2D layer with a kernel size of 2×2. The convolutional layers were followed by a flatten layer, a dense layer of 512 filters, a dropout layer with a dropout ratio of 0.2, and a final dense layer of 1 filter with sigmoid activation. The model was compiled with Adam optimizer, binary_crossentropy loss function, and accuracy metric. The training/validation data split was 5:5. The model was trained for 25 epochs, and its progress was monitored for validation loss. The training was rerun ten times, and the performance of the best models from each run was summarized.

The XGBoost classifier was trained in two ways. For the classification of original normal *versus* LUSC microarray data and LUAD *versus* LUSC RNA-seq data, the transcriptome was first filtered by removing the bottom 25% (quantile mean) genes and normalized by the Trimmed Mean of M-Values (TMM) method in edgeR. The training/validation split was 5:5, the random state was set at 7, and the learning rate was set at 0.3. For the classification of image-derived transcriptomes, the expression data was similarly filtered and normalized. The top 500 differentially expressed genes (DEGs) were identified from real transcriptome image-derived expression data by edgeR and validated with random forest in ranger with three-fold cross-validation. Fifty DEGs were selected as candidate hallmark genes according to the Gini coefficient ranking candidate gene importance in differentiating controls *versus* cases. The expression levels of 10 out of the top50 genes were too low (bottom 25%) in the GAN-synthesized LUAD and LUSC transcriptome images and not included in the final hallmark gene set. Therefore, the final classification model was trained with the 40 hallmark gene set with a training/validation split was 5:5, random state 7, and a learning rate of 0.3. The performance of the model was evaluated against the GAN-synthesized transcriptome data comprised of the 40 hallmark gene set.

The SRGAN discriminator consists of eight Conv2D layers (64×2, 128×2, 256×2, 512×2), and the generator has one Sequential block, six ResidualBlocks, and a UpsampleBlock. Two SRGAN models were trained: one for super-resolution from 512×512 to 1024×1024 pixels and another from 768×768 to 3072×3072 pixels. The first model was trained with 1840 paired images of 512×512 and 1024×1024 evenly sampled from 92 transcriptome classes from the TcgaTargetGtex dataset. LUSC transcriptomes from the TcgaTargetGtex were solely used for testing and not included in the training. The second SRGAN model was trained similarly with paired image libraries of 768×768 and 3072×3072 pixels. The 1024×1024 image produced by the first model was resized to 768×768 before feeding into the second model. The training was run at 7:3 train/validation split, crop_size 178, batch_size 4.

The StyleGAN2-ADA network consists of seven blocks (512-8) in the discriminator and eight blocks (4-512) in the generator network, and both have 65536 features. The conditional model was first trained with a 93 class, 46500 image library constructed from the TcgaTargetGtex dataset, then finetuned with a 93-class, 9300 image library re-sampled from the same TcgaTargetGtex dataset. After training, pseudo-LUAD and LUSC transcriptome images were synthesized with the generator from the conditional model.

### Statistical and functional enrichment analyses

Functional enrichment of genes in each of the compartments of the UMAP-transformed mouse protein-coding genome was performed at the DAVID web server.

Hierarchical cluster analysis of the overall subtypes of lung squamous cell carcinoma (LUSC) real and synthetic transcriptomes was performed in R using the hclust package with default settings using the transformed data scaled between [1, 255]. The differentially regulated genes between clusters were determined by a two-tailed *t*-test, and the top1000 genes with the lowest *p*-values were evaluated by DAVID for functional enrichment.

Median survival time (MST) analysis was performed in Prism 9. Genes with maximal variation across samples in each category were selected for analysis. The mitochondria electron transport chain complex I genes had lower variations across samples, and the average of the top six variable genes (NDUFA2, A6, B7, B11, S7, and V1) were taken.

### Data availability

All data needed to reproduce the results is available from GitHub (https://github.com/zjgt/Phoenix_Transformation).

### Code availability

All codes needed to reproduce the results is available from GitHub (https://github.com/zjgt/Phoenix_Transformation).

## Acknowledgment

I thank Angelo Pisco from Chan Zukerberg Biohub for technical assistance with the Tabula Muris Senis dataset. I thank Tero Karras, Derrick Schultz, Jun-Yan Zhu, and Lornatang for sharing their StyleGAN-ada, pix2pix, and SRGAN PyTorch repositories on GitHub, and Diego Porres, Janne Hellerstein, and Julian Pinzaru for technical assistance with the implementation. I thank Qiangfeng Zhang from Tsinghua for carefully reading this manuscript and for my past students’ hard work and inspiration. Guangzhou Laboratory supports this work.

---

**Supplementary Figure 1.**
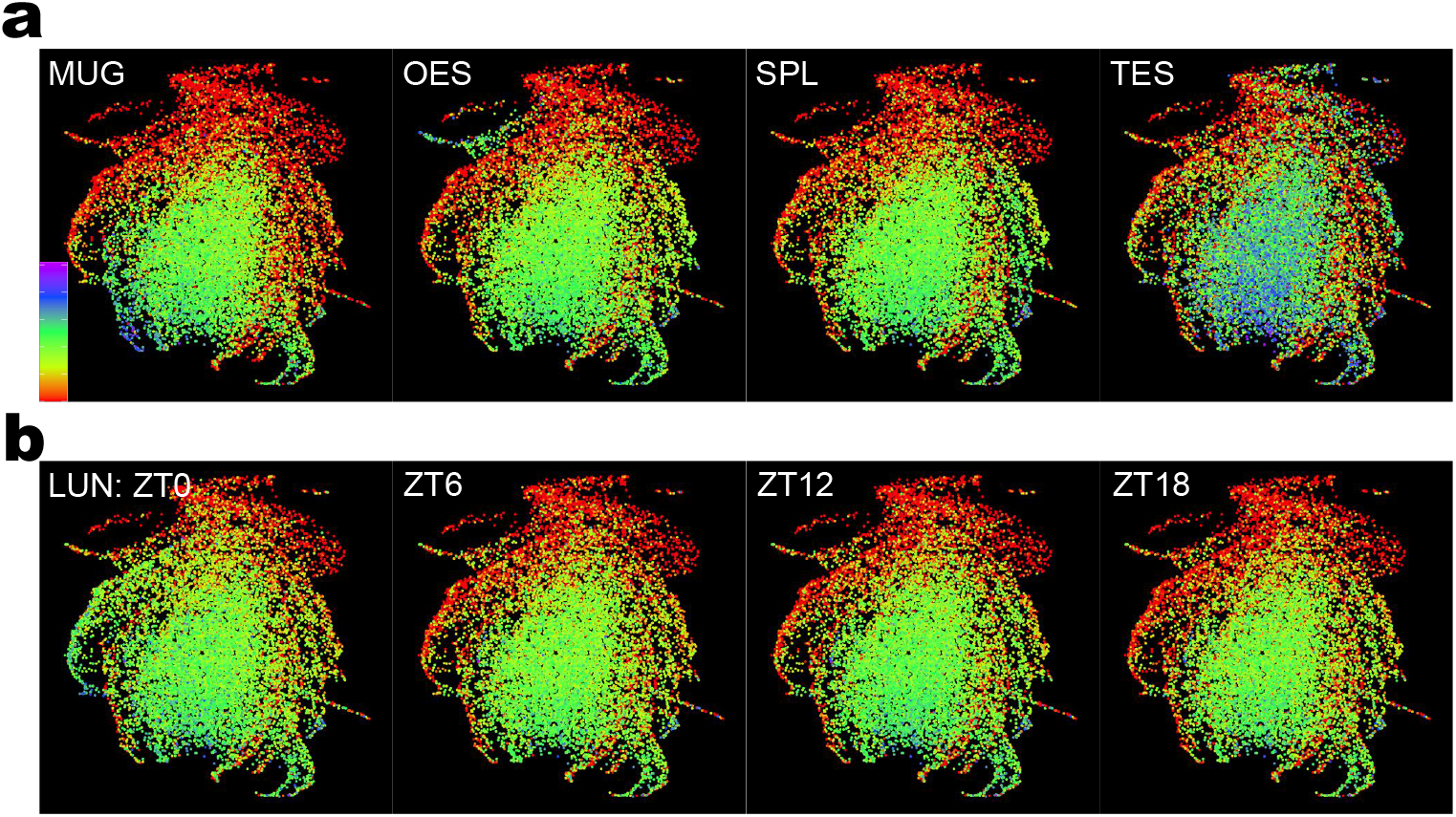
Baboon transcriptome visualization after phoenix transformation. **a**, Representative baboon tissue transcriptome images at ZT0. Color scale is set from 0 (red) to 255 (purple). Gene coordinates are transferred from mouse homologues. MUG: muscle gastrocnemius, OES: esophagus, SPL: spleen, TEST: testis. **b**, Representative lung (LUN) circadian transcriptome. Genes in the neuronal zone is specifically induced at ZT0. Color scale is the same as **b**. Gene expression data for transcriptome image synthesis are provided in Supplementary Table S8.

**Figure S2.**
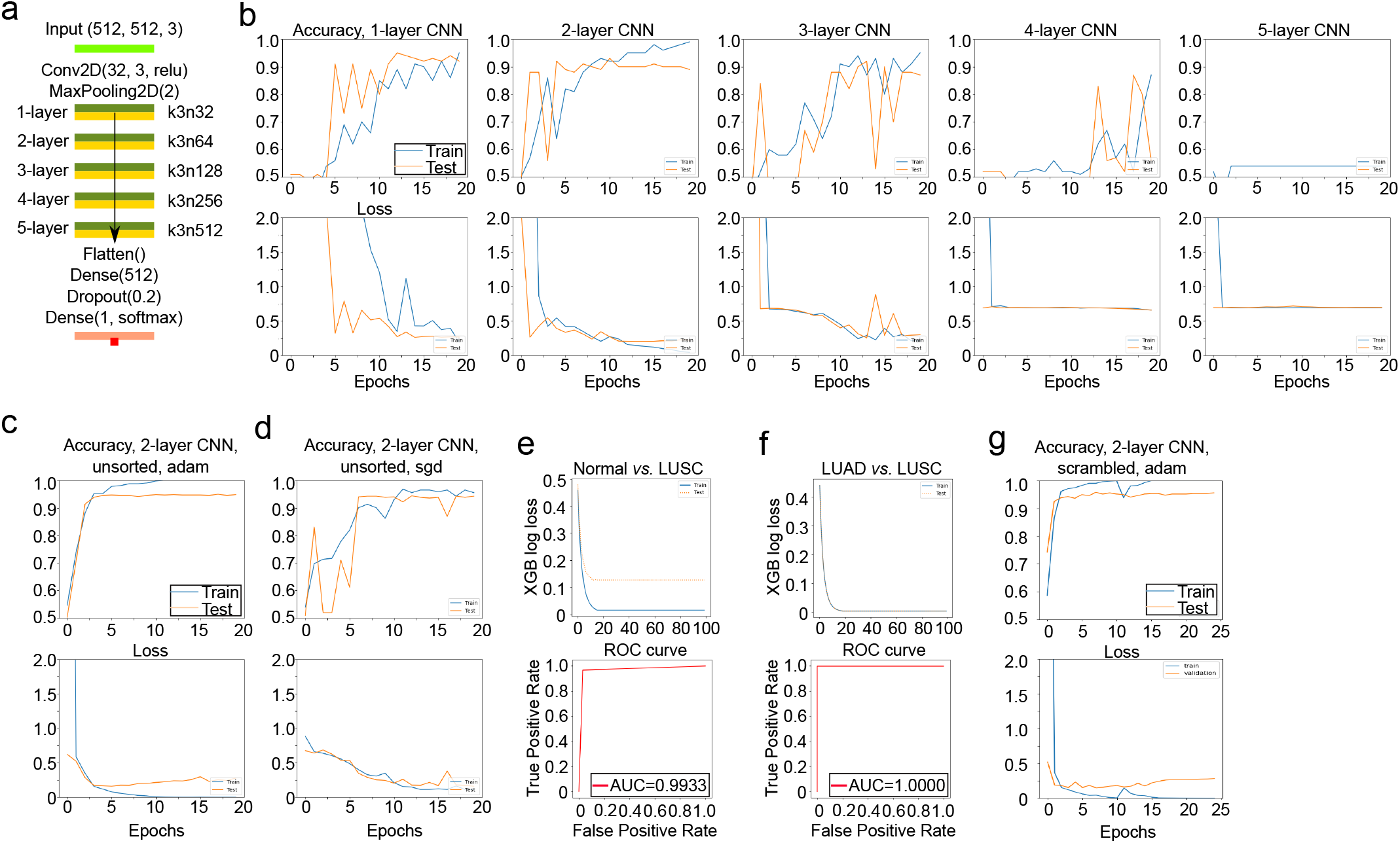
Convolutional neural network (CNN) parameter testing and benchmark with XGBClassifier. a, Illustration of a simple convolutional neural network with 1∼5 Conv2D layers and increasing filter sizes. b, Testing the performance of CNNs with 1∼5 Conv2D layers in LUSC detection with transcriptome images rendered in ascending order of gene expression and resized from 1024×1024 down to 512×512 pixels. **c-d**, Comparison the performance of adam (**c**) and stochastic gradient descent (sgd, **d**) optimizer in LUSC detection. **e-f**, LUSC detection (**e**) and LUAD *vs*. LUSC classification (**f**) by XGBClassifier with the same datasets. In both cases, LUSC is labeled as the positive state. Normalized gene expression data for LUSC vs. normal classification are provided in Supplementary Table S10, for LUSC vs. LUAD classification are provided in Supplementary Table S12. g, CNN model training with LUSC vs. LUAD transcriptome images where gene coordinates were scrambled.

**Figure S3.**
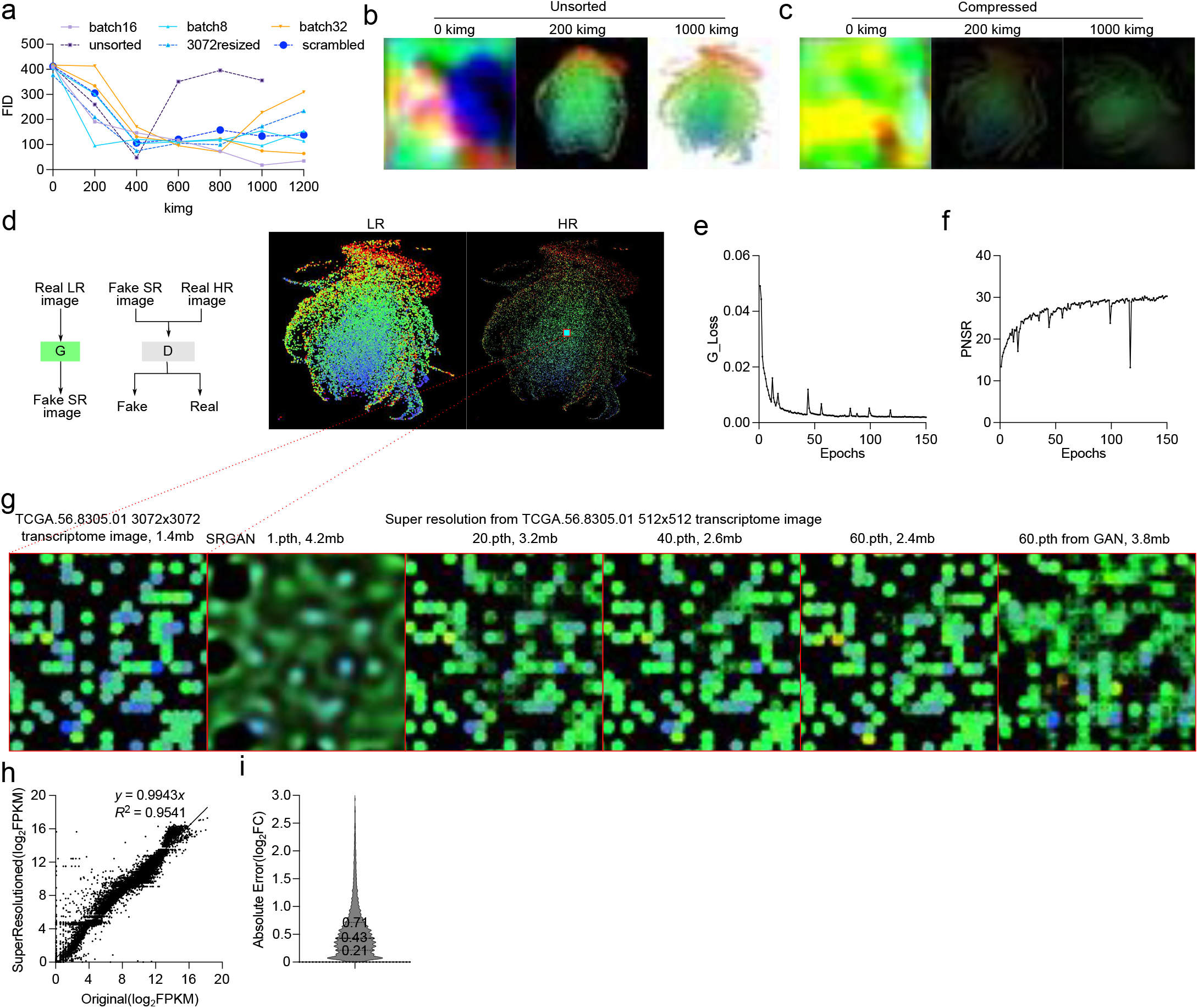
Generative adversarial network training for realistic transcriptome image and super resolution. **a**, Parameter testing for StyleGAN2-ADA training. Batch sizes of 8, 16, 32 were all trained with transcriptome images rendered in ascending orders of gene expression. Unsorted, images rendering without sorting; 3072resized, images resized from 3072×3072 to 512×512 pixels; scrambled, transcriptome images with gene coordinates scrambled. Traning was performed with default StyleGAN2-ADA parameters. **b-c**, Stalling of training using transcriptome images with either unsorted gene expression (**b**) or resized from 3072×3072 pixels (**c**). **d**, Illustration of the SRGAN network used for transcriptome image super resolution and gene expression information retrieval. The real LR image is 768×768 resized from 1024×1024, and the real HR is 3072×3072 transcriptome images generated from real transcriptomes in R. **e-f**, Training progress measured by G-loss (**e**) and peak signal to noise ration (PNSR, **f**) **g**, Illustration of the traning progress with images produced by the generator at different stages of the training process. The image file size is negatively correlated with image quality. The left most panel provides the ground truth high resolution image for the same segment as the super resolution image produced by the generator. **h-i**, Overall correlation (**h**) and mean absolute error (MAE,**i**) between the original expression data with gene expression information retrieved after a two-step super resolution. The original data are provided in Supplementary Table S14.

**Figure S4.**
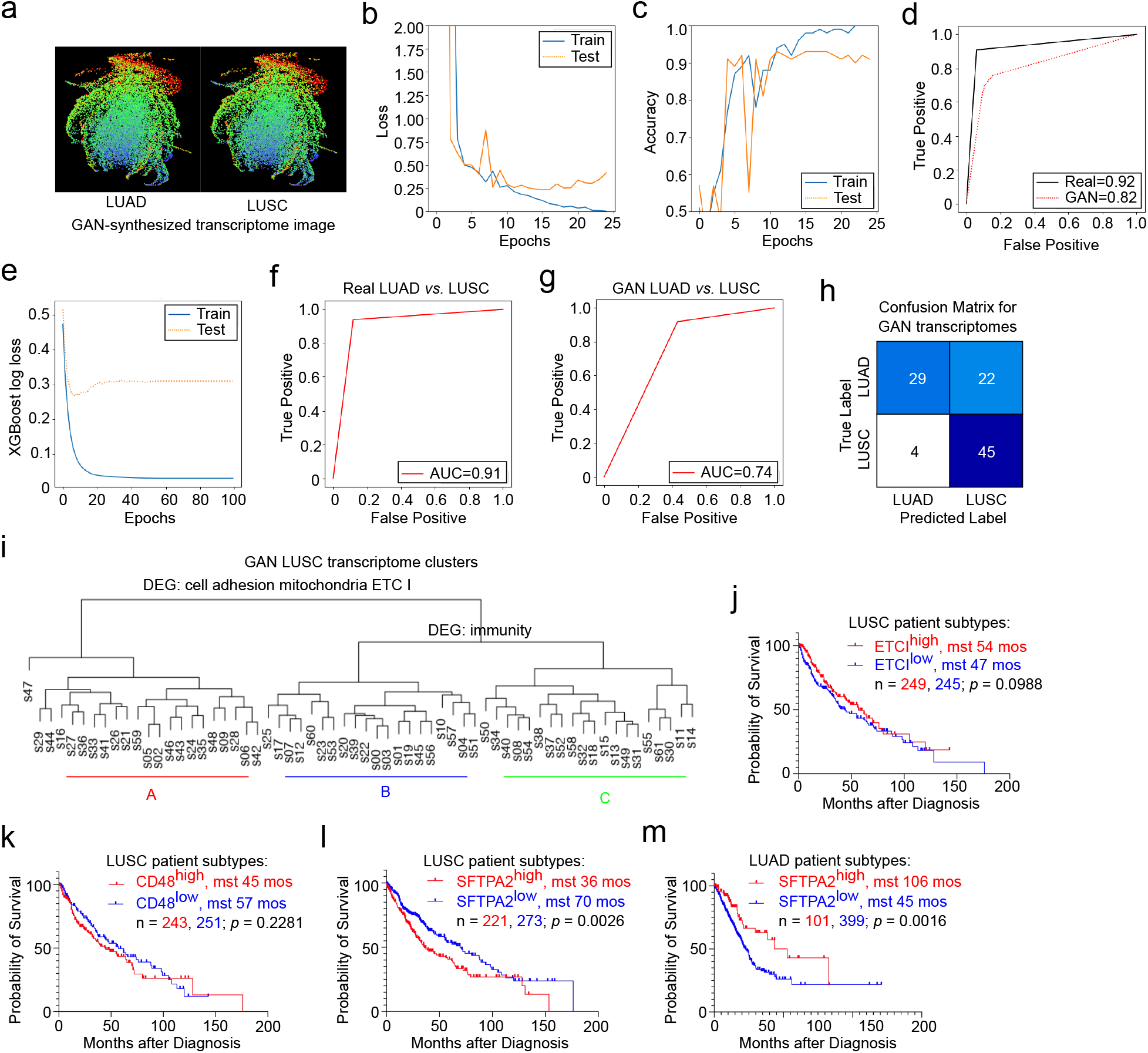
Synthesis and analysis of GAN-synthesized lung cancer transcriptome images. **a**, Realistic transscriptome images synthesized by trained StyleGAN2-ADA generative networks with seed 11, truncation 0.7, class=57 (LUAD) and 58(LUSC). **b-d**, Training of CNNs with real transcriptome-derived image (**b, c**) to classify GAN-synthesized LUAD vs. LUSC transcriptome images (**d**). **e-h**, Training of XGBClassifier with real transcriptome image-retrieved expression data (**e, f**) to classify GAN-synthesized LUAD vs. LUSC transcriptome image-derived gene expression profiles (**g, h**). 200 each of real and GAN-synthesized transcriptome images were super resolutioned to retrieve their gene expression signatures and analyzed. Gene expression data for classification are provided in Supplementary Table S23, 24. **i**, Eucaledian clustering of GAN-synthesized LUSC transcriptomes. A total of 62 LUSC transcriptome images were synthesized, super resolutioned to retrieve gene expression profiles, and clustered. Gene set enrichment analysis for real and synthetic LUSC subtyping is provided in Supplementary Table S25, 26. **j-m**, Kaplan-Meier plots of patient survival after dignosis with desginated gene expression signatures. The p-value was calculated from log-rank test for between-group survival difference.

